# Multi-group biodiversity theory and an application to multi-species maximum sustainable yield

**DOI:** 10.1101/2022.11.11.516177

**Authors:** A. Sadykov, K.D. Farnsworth, D. Sadykova

## Abstract

We introduce the group-based approach, use it to develop a multi-group biodiversity theory, and apply it find solutions to the multi-species maximum sustainable yield problem for a mixed species fishery. The group-based approach to community ecology is intermediate between classical species-centric and more recent trait-based (species-less) approaches. It describes ecological communities as composed of conspecific groups rather than species (as in classical models) or species-less individuals (as in trait-based models), and reconsiders community structure as results of inter-group resource competition. The approach respects species affiliation and recognises the importance of trait trade-offs at the conspecific group level. It offers an alternative to both classical and trait-based approaches and, remarkably, provides a complete analytical description of the community structure in the bench-mark case of zero-sum resource redistribution.

**Highlights:**

1. We introduce a group-based approach to modelling of ecological communities and develop a multi-group biodiversity theory.
2. A classification of intergroup interactions is established, based on the type of contest (which is determined by the relative role of qualitative vs. quantitative factors) and the accounting of resource redistribution.
3. For pure resource competition, we obtain the full analytic description of multi-group and multi-species community structure and its dynamics, including the processes of fission-fusion and invasion-extinction among groups.
4. A principle of competitive coexistence is formulated, which explains the existence of conspecific groups as a mechanism for avoiding competitive exclusion.
5. We apply the theory to harvesting multi-species communities (e.g., for fisheries) and derive analytic expression for total catch and approximate solutions for multi-species maximum sustainable yield (MSY).

## Introduction

Classical community ecology describes a hierarchy of interactions within ecological communities with three organizational levels: individual organisms, species, and the multispecies community. In this article, we develop an approach based on a different hierarchy: organisms, conspecific groups, and the multi-group community. Henceforward, by conspecific group (briefly, a group), we mean a stable self-organized group of organisms of the same species living together and engaging in some form of group behaviour (e.g., information exchange, forging, defence, and reproduction). (Note we use ‘group’ only to refer to a set of organisms, never in the mathematical ‘group theory’ sense). Essentially, conspecific group is an umbrella term that generalises various species-specific terms such as pack, pride, flock, family, colony, herd, swarm, shoal, etc. (Krause and Ruxton 2002). It is distinct from the concept of meta-population as the group is not necessarily isolated by environment, or genetically: if a local population L, composed of individuals, is represented as a member of a meta-population M, then the group G is a subset of M, where M contains at least one group. A conspecific group is a set of individuals whose fitness is partly determined by the fitness of other members of their group (Haldane 1932). In cases of obligatory group behaviour, it is relatively easy to identify groups. However, in other cases, a detailed analysis of inclusive fitness and genetic relatedness may be required to establish a conspecific group structure within a local population. In the case of microbiological communities, where the definition of a species can be unclear (due to rapid evolution and horizontal gene transfer), the group-based (or strain-based) approach may be the most appropriate choice for population dynamic models.

The classical approach using a single population to represent each species overlooks the structure of intraspecific competition, which may consist of two distinct types of interactions: within a group and among groups. When intra-group interactions are more cooperative in nature (Hamilton 1963), then usually inter-group interactions show fierce competition. The nature and structure of intraspecific competition can determine the conditions for species coexistence and, accordingly, community assemblage and biodiversity, all of which is lost in the classical approach.

The group-based approach considers intraspecific interactions by recasting the ecological community as a multi-group community (i.e., set of interacting conspecific groups). This does not imply a rejection of the species-based approach, but rather reconsiders the species population as a metapopulation of conspecific (single-species) groups whose numbers are established through a process of inter-group competition. For this reason, the group approach is more consistent with empirical field ecology where observations are typically made at the scale of individuals and groups of organisms, then scaled up to area and species levels, post-hoc.

Within a multispecies community, species are typically performed multiple ecological functions, making it difficult (sometimes impossible) to assign a single role (predator or prey, resource, or consumer, etc.). For example, most fish species are both predators and prey for each other, depending on body size (for which size-structured trait-based models have been developed by Andersen et.al. 2016, Kiørboe et.al. 2018). To cope with these difficulties, we apply the concept of the trophic chain, i.e., a consumer can feed directly on autotrophs and/or species that feed on autotrophs and/or species that feed on species that feed on autotrophs, and so on. In such a food chain, the diet of each consumer can be step by step traced back to the autotrophic level. It allows to recalculate the diet of each consumer in terms of the consumption of basic (autotrophic) resources and reconsider the food web as a resource-consumer system, where the common resource is represented by autotrophic species, and all other species are considered as direct or indirect consumers (i.e., predators). Generally, it allows one to redefine the outcome of various ecological interactions in terms of competition for resources with additional predation mortality and to model ecological communities based on conspecific groups competing for a common resource. In the first section, we introduce the multi-group resource competition model and identify three (positive-sum, negative-sum and zero-sum) generic cases of the system. In the second section, we analyse the multi-group system at different set of parameters and formulate the competitive coexistence principle. In the third section, we develop the multi-group biodiversity theory for the zero-sum case. In the fourth section, we describe structure of multi-group community. In the fifth section, we consider fission-fusion and invasion-extinction dynamics within multi-species community. In the sixth section, we apply the multi-group theory to the multi-species maximum sustainable yield problem. Finally, we discuss the similarities and differences between multi-group and other biodiversity theories.

### 1. Resource competition model

The group-based approach models competition for resources at two levels: among groups and within groups. Within group competition is described in Sadykov 2011; here we concentrate on competition among groups, since at the community level, the resource-based group population dynamics can be written in general as

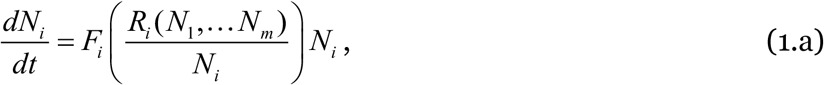

where *N_i_* is the population size and *F*(*R_i_*(*N*_1_,…*N_m_*)/*N_i_*) is the net growth rate of i-th group. The function *F_i_* depends on a species and group specific set of density-independent parameters (they are omitted here) and on the density-dependent average resource for a given group *R_i_*(*N*_1_,…*N_m_*)/*N_i_*, where *R_i_* are resource share of i-th group. Here it is not necessary to specify the form of the function *F_i_*(*N*_1_,…*N_m_*), it must satisfy the standard requirements of smoothness and reversibility, and may incorporate several density-dependent processes, including predation.

At the inter-group level, the resources sharing process can described by generalized Tullock contest function, which is widely used in economics (Tullock 1980, Konrad 2009) and evolutionary biology (Hawkes et. al. 1995, Kokko and Morrell 2005, Reeve and Hölldobler 2007, Crowley and Baik 2010, Gavrilets 2012).

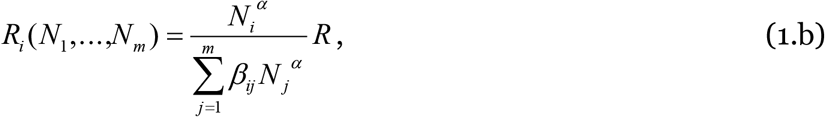

where *R* is total available at some time interval resource, *β_ij_* are coefficients that describe the per-capita effect of the j-th group on the i-th group share and *α* is a parameter, that quantifies the sensitivity of resource competition outcomes to group sizes. At *α* = 0, the share of resources obtained by the group does not depend on its size, at *α* →∞, the largest group takes all the resources. Equation 1b expresses a very general model of a *competitive milieu* in which quantitative, density-dependent features (i.e., group sizes) combine with qualitative, density-independent characteristics (i.e., the relative percapita influence of competitors on one another).

When the resource rate *R* is constant the system of equations (1.a and 1.b) describes resource competition between groups. The solutions of the equations x.1a for fixed points (*N* = 0) can be written as

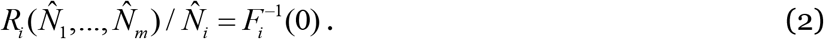

In the special a single group, this solution can be rewritten as 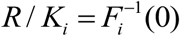, where 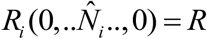 and 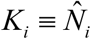, which can be called a *singular carrying capacity* for the i-th group (i.e., the stable population size of the i-th group in the absence of other competitors). Noticing that the right side of the equation 2 is a constant, one can obtain the basic principle of multi-group community dynamics:

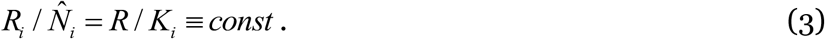

which can be called the principle of preservation of the average consumption rate and states that *in a multi-group community, the equilibrium population size of each group is such that the average resources per individual remains constant*. In other words, the group responds to changes in the realised resource (i.e., obtained in competition) by changing its population size, while maintaining the average level of consumption.

Now we can establish the formal connection between the systems of multi-group dynamics equations (1.a, 1.b) and the competitive Lotka-Volterra equations (Townsend et al. 2008). Substitution of equation 3 into the equation 1.b, with some re-arrangement gives:

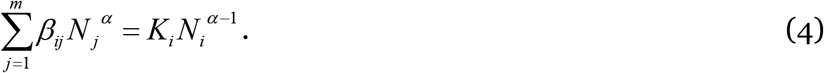

If we further, suppose that all diagonal elements *β_ij_* are equal to 1 and rewrite the equation x.4 we get a system

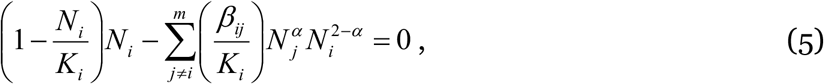

which can be considered as a system of zero-growth isoclinic (nullcline) equations for the following system of differential equations:

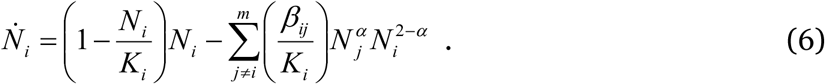

The systems of equations 1 and 6 are analogous (isomorphous) in the sense that they have the same nullclines, meaning they have the same fixed points and conditions of stability, although specific solutions for these systems may differ. Note that at *α* = 1, the system 6 is identical to the competitive Lot-ka-Volterra equations.

Resource sharing equation x.1b gives us some insight on the community matrix **β**_ij_. In the general case, Σ *R_i_* is not necessary equals to *R*, with the effect that additional resources Δ*R* = Σ *R_i_* – *R* may appear (Δ*R* > 0) or disappear (Δ*R* < 0). The former can be regarded as a “positive-sum game” or competition with additional cooperative effect, with the latter a “negative-sum game” or competition with additional (interference) adverse effect. Later, we will discuss these cases, but now let us consider the case of a zero-sum game i.e., pure resource competition for which Δ*R* = 0. If we suppose that *β_ij_* coefficients are

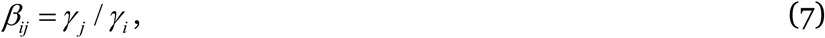

then we can rewrite equation (1.b) as

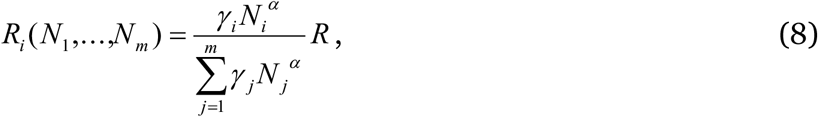

which obliviously satisfies the Δ*R* = 0 condition. Conditions 7 collapse the community matrix **β**_ij_ into the vector *γ*_i_, thus all the variety of influences of the i-th group on other groups is reduced to one value of *γ_i_*, which can be called the *competitive strength* of the i-th group, since this value accumulates all the qualitative, density-independent factors of resource competition. It broadly incorporates all the individual traits of a member of the i-th group that determine the result of inter-group resource competition. Also, conditions 7 can be called a transitivity condition since it implies that if *γ_i_* > *γ_k_* and *γ_k_* > *γ_j_* then *γ_i_* > *γ_j_*. It is important to emphasize that a single measure of competitive strength exists only in the case of a “zero-sum game”, in other cases, where the transitivity condition is not met, it is impossible to unambiguously determine which of the competing groups within the community is stronger or weaker. Another important consequence is that an explicit closed resource competition model (one that satisfies the conservation law) exists only if the transitivity condition is satisfied. For example, any resource competition model that combines resource dynamics with Lotka-Volterra type of population dynamics (which is a positive sum game in the case of species coexistence) cannot be closed unless it explicitly includes an extra equation for the additional resource dynamics Δ*R*(*t*).

### 2. Competitive coexistence principle

Up to now the competitive exclusion principle (Hardin 1960) or Gause’s law remains a milestone in ecology despite multiple controversies (Pocheville 2015) and paradoxes (Hutchinson 1961). A common feature of these debates is that they consider the competitive exclusion and coexistence as the result of interactions between species, although Gause himself formulated the problem in the following way: *“the struggle for existence among animals is a problem of the relationships between the components in mixed growing groups of individuals, and ought to be studied from the viewpoint of the movement of these groups.”* (Gause 1934). In other words, Gause saw the solution of the coexistence problem not in interactions between species, but rather in interactions between groups of individuals. In this section we investigate how the group-based perspective extends the species-based views of competition and coexistence.

As shown in the previous section, the main difference between the group-based competition model and the classical Lotka-Volterra one is the inclusion of the *α* parameter, which generalises competitive effects. We interpreted *α* as the type of contest: a measure of the balance between the quantitative (group population sizes) and qualitative (relative competitive strengths) factors in inter-group resource competition (Table 1).The system of equations x.6 enables us to reinterpret the parameter *α* in two other ways, which are more widely used in ecology (Table 2). First, we can consider *α* as a parameter that determines the balance between self-regulation (first term in equation 6) and exogenous regulation (second term in equation 6). If *α* is less than 1, the first term is always greater than the second at sufficiently small population size. This means that whatever the exogenous pressure, the population always has the potential to avoid extinction by reducing its numbers. Under such conditions, any number of groups competing for one resource can coexist (this mechanism can be called as *“refuge at small numbers”)*. Note that here we describe the general mechanism in a mathematical sense, in practice a sufficiently small population size should exceed a viable population size.

**Table 1.** Description of different types of contests in relation to the value of α and resource competition outcome.

| Value of $\alpha$ | Contest type | Expected result of resource competition | |
| --- | --- | --- | --- |
| $\alpha \rightarrow \infty$ | **“Winner-takes-all”**, the most numerous group grabs all resources regardless its relative competitive strength | Conditional exclusion | $N(0,..N_j,..,0)$, $N_j=K_j$, where j depends on initial group sizes |
| $\alpha > 1$ | **“Quantity over quality”**, the resource share of each group depends more on its numbers than on competitive strength | Conditional exclusion | $N(0,..N_j,..,0)$, $N_j=K_j$, where j depends on initial group sizes |
| $\alpha = 1$ | **“Quantity – quality balance”**, numbers and relative competitive strengths equally important for resource acquisition | Competitive exclusion | $N(0,..N_j,..,0)$, $N_j=K_j$, where j is defined by $\text{Max}(\gamma_j K_j)$ |
| $0 < \alpha < 1$ | **“Quality over quantity”**, the resource share of each group depends more on its competitive strength than on numbers | Coexistence | $N(N_1,..N_i,..,N_m)$, $N_i=CP_i K_i$, where $CP_i$ is the relative competitive power |
| $\alpha \rightarrow 0$ | **“Only quality matters”**, the resource is distributed according to the competitive strength regardless of the size of each group | Coexistence | $N(N_1,..N_i,..,N_m)$, $N_i=CS_i K_i$, where $CS_i$ is the relative competitive strength |

**Table 2.**
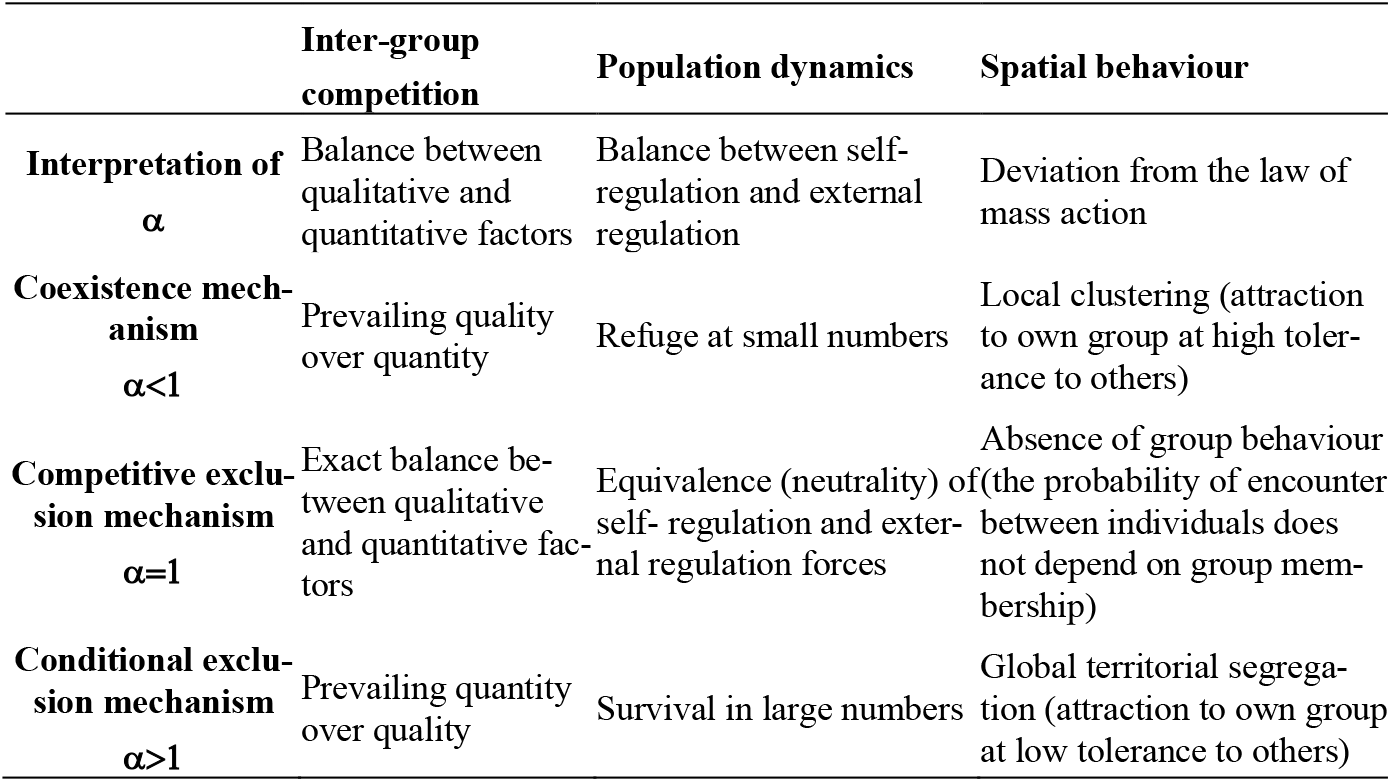
Three equivalent interpretations of a and mechanisms of competition from the point of “inter-group competition”, “population dynamics” and “spatial behaviour” perspectives. Note: in the classical ecology for the “self-regulation” and “external regulation”, the terms intraspecific and interspecific are usually used. We avoid doing this because inter-group competition can be between different groups of the same species as well as between groups of different species organisms.

In the opposite case at *α* > 1, the external regulation dominates, and the net growth rate is positive only for sufficiently large populations (fig.1). This implies that the group can survive only with sufficiently large initial population (this mechanism of conditional exclusion can be called *“survival in large numbers”)*.

**Fig.1.**
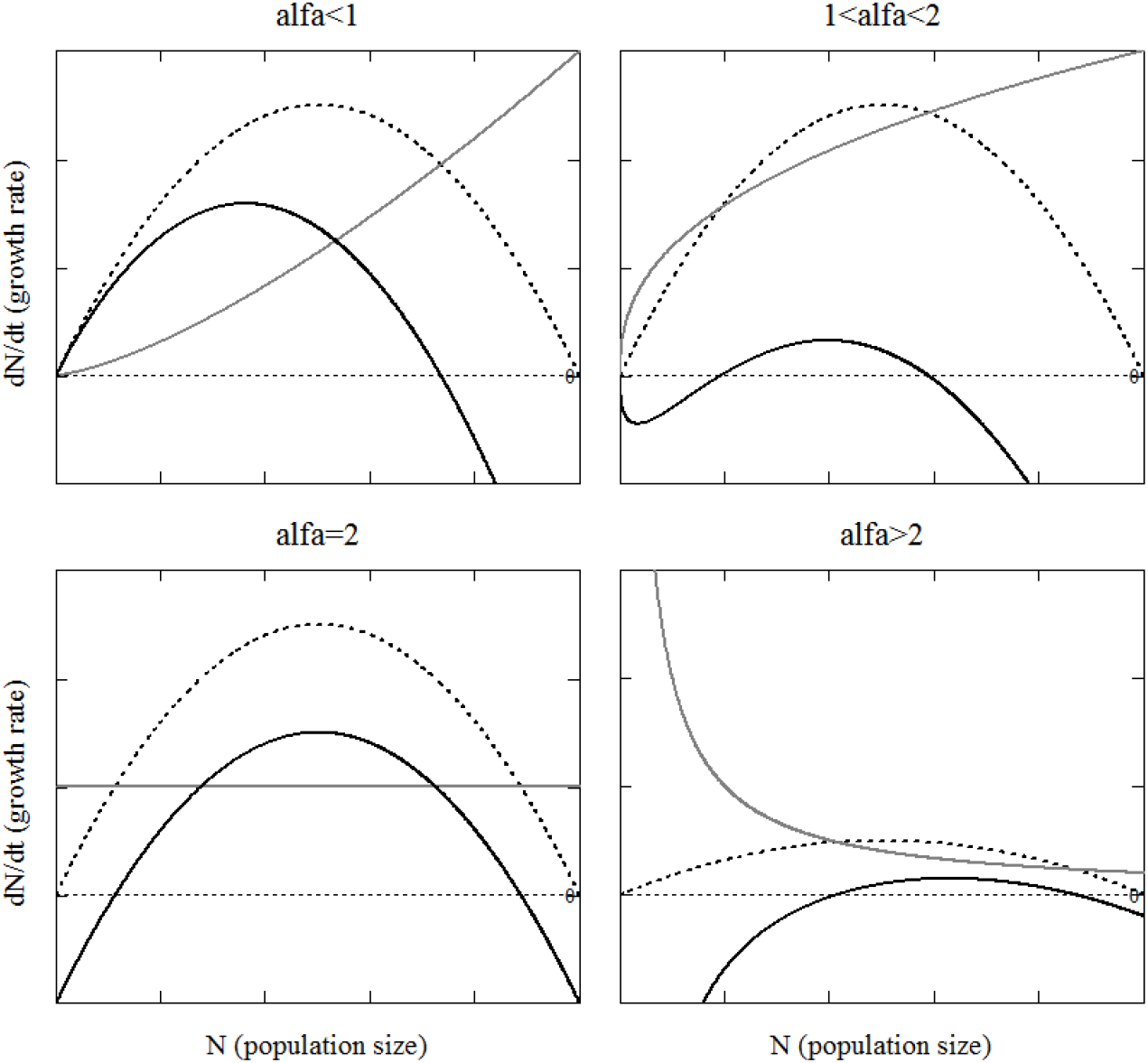
The net growth rate (black bold line), the self-regulation term (black dotted line) and the external-regulation term (grey line) at different value of α.

Secondly, we may interpret the *α* as a parameter that describes the relationship between the probabilities of encounters between groupmates and members of other groups. If we ignore group membership, then the second (interaction) term in equation 6 is proportional to the square of the total number, as it should be for a well-mixed (randomly distributed) in space population. However, if we consider group membership, we note that the divergence of *α* from 1 determines the deviation from the expected (for well-mixed populations) law of mass action. If *α* < 1, then the interaction term is less than would be expected from the law of mass action at small numbers and greater than expected for large numbers. This suggests *local clustering* in the spatial distribution of competitors. In the case *α* > 1,we see the converse; the interaction term is greater than expected for small numbers and less than expected for large numbers. That dependence is characteristic of the boundary-to-area relation, which would indicate *global territorial segregation*. These spatial relationships resemble the Schelling’s model of segregation (Schelling 1978, Vinković and Kirman 2006), in which both local clustering and global segregation emerge for high and low tolerance to neighbours, respectively.

Next, we describe the role of the matrix **β**_ij_. As mentioned in the first section, the elements of this matrix determine the overall type of competition with respect to its effect on total realised resource 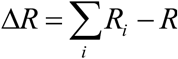. There are three generic cases:

1. Δ*R* < 0, the negative-sum game in which total realised resource is less than the resource present (2) Δ*R* > 0, the positive-sum game in which realised resources is greater and (3) Δ*R* = 0, the zero-sum game in which total realised resource is the same. Note that Δ*R* → (*m* – 1)*R* at *β_ij_* → 0 and Δ*R* →-*R* at *β_ij_* → ∞, which indicates that “weak interactions” lead to the positive-sum game, while “strong interaction” to the negative-sum case. In the zero-sum case, “strong” and “weak” interactions are balanced within each pair of competitors, since *β_ij_* = *γ_j_* / *γ_i_* implies *β_ij_β_ij_* = 1. For the elements of the matrix **β**_ij_, there is a value 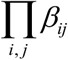, which can be used as an indicator of deviations from this balance of interactions. 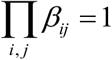 responds to Δ*R* = 0 (i.e., the zero-sum case), 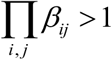 to Δ*R* < 0 (i.e., the negative-sum case), and 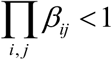 to Δ*R* > 0(i.e., the positive-sum case). It should be noted that this condition is necessary but not sufficient and does not cover all possible cases, for example, when one or more *β_ij_* are equal to zero.

We can now summarise inter-group competition, based on the values of *α* and **β**_ij_ together (Table 3). In five of the nine cases, inter-group competition can lead to coexistence, but the mechanisms behind each are quite different. Let us consider them in detail.

**Table 3.** The inter-group competition outcomes at different ranges of α and |β_ij_|. The middle column (α=1) may refer to the Lotka-Volterra competition model, while the middle row refers to the pure (zero-sum) resource competition examples.

| | $\alpha < 1$ | $\alpha = 1$ | $\alpha > 1$ |
| --- | --- | --- | --- |
| Resource competition with<br>adverse effect $\Delta R < 0$ (neg-<br>ative-sum game)<br>$\Pi\beta_{ij} > 1$ | Coexistence of<br>less than MG groups | Conditional exclusion if<br>$\beta_{ij} \beta_{ji} > 1$, $\beta_{ij} > K_i/K_j$ and<br>$\beta_{ji} > K_j/K_i$, otherwise<br>Competitive exclusion | Conditional<br>exclusion |
| Pure resource competition<br>$\Delta R = 0$ (zero-sum game)<br>$\Pi\beta_{ij} = 1$ | Coexistence of<br>up to MG groups | Competitive exclusion | Conditional<br>exclusion |
| Resource competition with<br>cooperative effect $\Delta R > 0$<br>(positive-sum game)<br>$\Pi\beta_{ij} < 1$ | Coexistence of more<br>than MG groups | Competitive exclusion or<br>Coexistence if $\beta_{ij} \beta_{ji} < 1$,<br>$\beta_{ij} < K_i/K_j$ and $\beta_{ji} < K_j/K_i$ | Conditional<br>exclusion<br>or Coexistence |

When *α* < 1 (the first column in the table 3), coexistence is possible because at a small enough population, potentially, any group has a positive growth rate. However, this does not mean that any number of groups can coexist, since their number is limited by available resources. It should be noted that coexistence in this case is *evolutionarily stable*, because possible mutations (including those that affect the physiology of individuals) does not lead directly to the extinction of any groups, but rather to changes in their relative sizes.

When *α* = 1 (the Lotka-Volterra case), coexistence is possible if and only if the realised resource greater than *R*. In theoretical ecology the implied “resource surplus” is usually explained in one of two ways: implicitly through *niche separation* (MacArthur and Levins 1967), where the total resource available for competitors is greater than the resource over which competition is taking place or explicitly by introducing several types of resources (Tilman 1977). However, in contrast to the previous case, coexistence here is *evolutionary unstable*, because even for a few competitors, stable coexistence is possible only within a very narrow range of parameters, so even small changes in the phenotypes can directly lead to the extinction of most of the rival groups (May 2019).

Finally, for *α*> 1, coexistence is also possible if the realised resource is larger than each group would receive in the absence of competition. Such circumstances could only occur when the potential benefits of cooperation outweigh the possible losses from competition. However, as in the previous case, coexistence here requires very precise coordination of parameters likely disrupted by mutations and further, *α* > 1 implies a strong spatial segregation of groups (as an aside, we could interpret competition for resources among cells within an organism with this formulation).

Now we can formulate the principle of competitive coexistence for inter-group competition: *A significant number of (irrespective of whether single or multi species) groups competing for the same limited resource can coexist as an evolutionary stable system, if they are involved in group behaviour and the qualitative factors of competition prevail over the quantitative*. It should be noted that this principle includes the competitive exclusion principle as a special case for 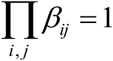 and *α* = 1 (the middle cell in table 3), which can be rephrased as follows: “pure (zero-sum game) resource competition in the absence of group behaviour leads to competitive exclusion”. In the limiting case of a single species, the principle of competitive coexistence reduces to the assertion that several groups can coexist sympartically, which is consistent with empirical observations. Accordingly, the principal provides a fundamental ecological explanation for the existence of groups among competing organisms: they form groups, because this allows them to avoid competitive exclusion.

In general, the group-based approach allows us to bypass specific for the species-based approach limitations, such as the paradox of plankton (Hutchinson 1961) and develop a more general theory of biodiversity based on inter-group interactions.

### 3. Multi-group theory of biodiversity

In this section, we investigate the multi-group community in the case where 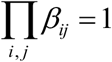 and *α* < 1 (the zero-sum case), for which the exact analytic solution for fixed point 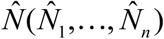 can be obtained as

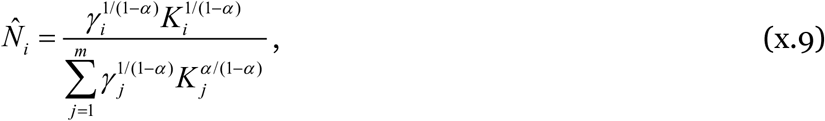

that can be verified by direct substitution into equation x.8. This nontrivial (all populations are non-zero) fixed point is a stationery focus at *α* < 1 or a saddle point at *α* > 1. The former means that all groups can coexist, while later means there is a *conditional exclusion* i.e., only one group survives, depending on the initial conditions (population sizes). At *α* → 1, the term with the maximum *γ_j_ K_j_*. dominates and the fixed point tends to 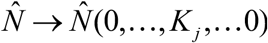. In other words, at *α*→ 1 a *competitive exclusion* occurs, i.e., only one group with the largest *γ_j_ K_j_* survives.

Also, one can rearrange equation 3 as 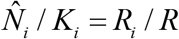, then take the sum of both sides of this equation given equation 8 and get 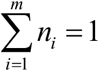, where *n_i_* ≡ *N_i_*/*K_i_* represents the resistance to competitive pressure which is the reciprocal of the niche contraction coefficient (Carvalho and Cardoso, 2020) since in ecological terms 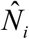 and *K_i_* are the realized (post-competitive) and fundamental (pre-competitive) niche sizes, respectively. Thus, one more principle can be formulated: *in a multi-group community, under the condition of pure resource competition, the total sum of inverse niche contraction coefficients is always equal to one*. In essence, this principle is a reformulation of the conservation law for the ecological community, a consequence of the assumption that all available resource is, in some way, divided among groups and consumed by individuals.

For convenience, the following substitution of variables can be introduced: *α* ≡ *D*/(*D* +1) and *C_i_* ≡ *γ_i_K_i_*. The quantity *D* can be called a *competitive exclusion index* since an increase in this index means approaching the competitive exclusion limit (i.e., 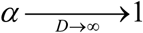). The quantity *C_i_* can be called a *competitive ability* of the i-th group, since in the competitive exclusion limit only the one group with maximum competitive ability survives. With these preconditions, the equations for equilibrium population of the i-th group 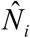, the total population of the multi-group group *N*_Σ_, the relative abundance 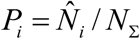, and the inverse niche contraction coefficient 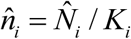 can be written as

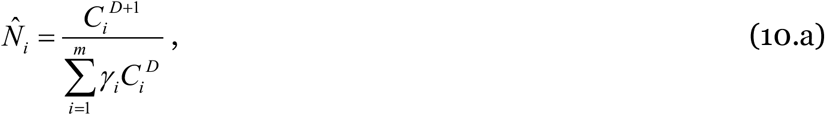

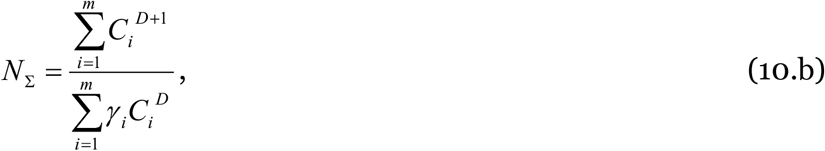

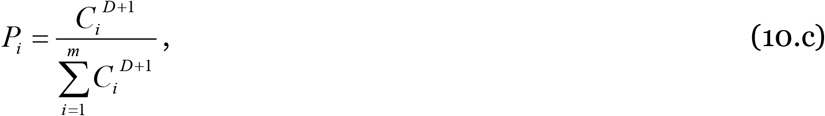

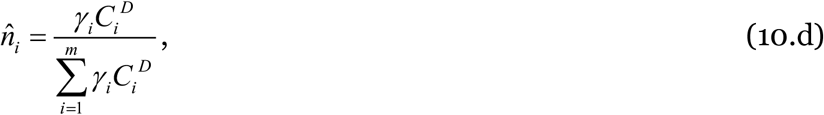

For further analysis, we decompose equation x.10b into expected values *E*[*X*] and covariance between variables cov(*X, Y*) in several ways as follows:

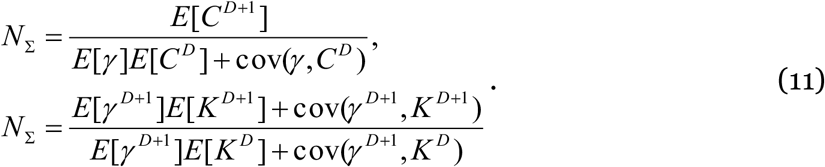

Since covariance with a constant is zero, we can obtain the total population size of the community for some special cases (Table 4).

**Table 4.** The i-th group population, the total population size and diversity (effective number of groups) of the multi-group community in some particular cases.

| | $N_i$ | $N_\Sigma$ | $GD$ |
| --- | --- | --- | --- |
| Neutral case 1,<br>All competitive abilities<br>are equal<br>$\forall \gamma_i K_i = C$ | $\frac{C}{\sum_i \gamma_i}$ | $m \frac{C}{\sum_i \gamma_i}$ | $m$ |
| Neutral case 2,<br>All competitive abilities<br>and all $\gamma_i$ are equal<br>$\forall \gamma_i K_i = C, \forall \gamma_j = \gamma$ | $\frac{C}{m\gamma}$ | $\frac{C}{\gamma}$ | $m$ |
| All competitive<br>strengths are equal<br>$\forall \gamma_j = \gamma$ | $\frac{K_i^{D+1}}{\sum_{i=1}^m K_i^D}$ | $\frac{\sum_{i=1}^m K_i^{D+1}}{\sum_{i=1}^m K_i^D}$ | $\frac{\left(\sum_{i=1}^m K_i^D\right)^{D+1}}{\left(\sum_{i=1}^m K_i^{D+1}\right)^D}$ |
| All currying<br>capacities are equal<br>$\forall K_j = K$ | $\frac{\gamma_i^{D+1}}{\sum_{i=1}^m \gamma_i^{D+1}} K$ | $K$ | $\frac{\left(\sum_{i=1}^m \gamma_i^D\right)^{D+1}}{\left(\sum_{i=1}^m \gamma_i^{D+1}\right)^D}$ |
| Both all currying capaci-<br>ties and all competitive<br>strengths are equal<br>$\forall \gamma_j = \gamma$<br>$\forall K_j = K$ | $\frac{K}{m}$ | $K$ | $m$ |
| Competitive exclusion<br>limit<br>$D \rightarrow \infty$ | $\frac{\max(K_i \gamma_i)}{\gamma_i}$ | $K_i$ | 1 |
| “Only quality matters”<br>type of contest limit<br>$D \rightarrow 0$ | $\frac{\gamma_i K_i}{\sum_{i=1}^m \gamma_i}$ | $\frac{\sum_{i=1}^m \gamma_i K_i}{\sum_{i=1}^m \gamma_i}$ | $m$ |

Using the relative abundances (10.c) we can calculate *a group diversity index GD*, which is a special case of the *true diversity index* (Hill 1973, Jost 2006), 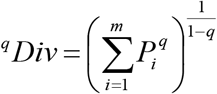 with *q* = *α*= *D*/(*D* +1) as:

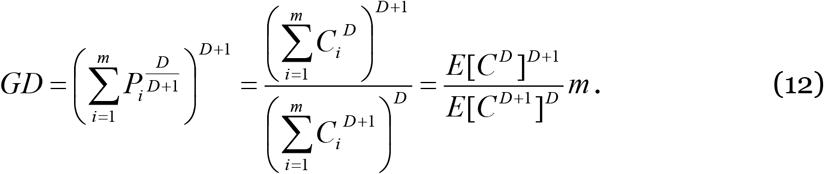

This index *GD* ≤ *m* represents an effective number of groups in the multi-group community (i.e., members of these groups most regularly appear in samples from this community). Values of *α* less than 1 mean that rare groups are detected more frequently than would be expected based on their proportional abundances alone. Based on this index, *the principle of soft competitive exclusion* can be formulated: in a community potentially consisting of *m* groups, only members of *GD* groups can be regularly sampled. Note that in the limit *D* →∞ this principle converges to the principle of competitive exclusion (Table 4). According to equation 12, there are two reasons for declining the diversity of the community: (1) increasing the role of quantitative (relative to qualitative) factors for competition (*D →∞*) and (2) deviation from the neutrality condition (∀*C_i_* ≡ *const*), i.e., increasing of inequality in the competitive abilities between groups. Significantly, this index can be directly estimated from empirical data, which opens the way for a detailed verification of the multi-group theory. Along with the group diversity index *GD*, another *competition diversity index GN* can be introduced as:

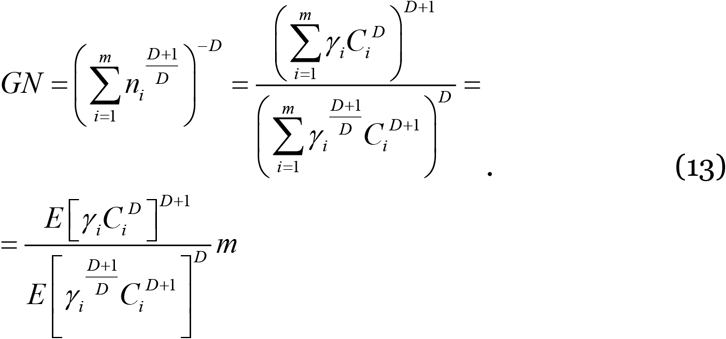

This index estimates the effective number of groups, which are resistant to competitive pressure (i.e., whose fundamental niche is less contracted) in the multi-group community. A combination of these indices can be useful for describing and monitoring community dynamics in term such as normal turnover *m* – *GD* (expected number of rare species/groups in the sample) and invasion turnover *m* – *GN* (i.e., turnover above this value indicates a change in the structure of competition in the community).

As a demonstration of the efficiency of this theory (i.e., the ability to produce empirically verifiable predictions based on first principles, using a few free parameters, (Marque *et al*. 2014), we will derive the relationship between diversity and total community population. Assuming that *C*~ *Lognorm*(*μ,σ*^2^) is log normally distributed, using the equations 10.c, 11 and 12 one obtains:

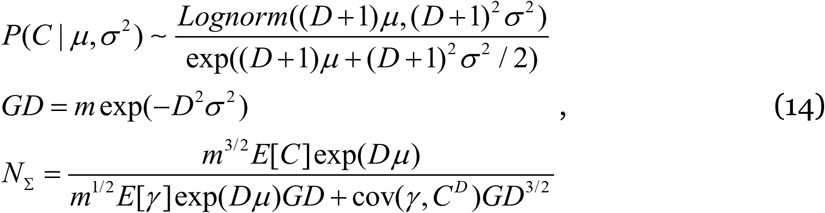

These equations show several things. Firstly, diversity declines as the system moves away from neutrality (i.e., increasing *σ*) or/and approaching the competitive exclusion limit. Secondly, the size of the species/groups regional pool *m* positively affects its abundance. Thirdly, in the case of a positive correlation between quantitative (i.e., *K*) and qualitative (i.e., *γ*) factors of competitive abilities, the total abundance declines as diversity increases. Finally, in the case of a negative correlation, there is an intermediate level of diversity 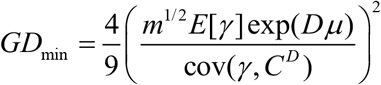 at which total abundance is minimal, i.e., both low and high diversity can support elevated community abundance.

Although the results above are presented in terms of group populations it is possible to rearrange all of them in terms of species populations by summing over groups of the same species 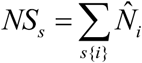, where *s*{*i*} is a subset of group indices related to a particular species.

### 4. Population structure of multi-group communities

From formulas 10.a and 10.b, it is clear that changes in the singular carrying capacity and/or competitive strength of a particular group affect the populations of other groups and hence the population of the multi-group community as a whole. Therefore, it is possible to classify each group according to the degree of its influence it has on the multi-group community and thus reveal the structure of this community. To test this intuition, we calculate the values of some partial derivatives (Table 5) that can be considered as the sensitivity of population in one group to the properties of another group (or the entire community). In the last two rows of Table 5, we see that the total population reacts differently to changes in groups depending on their properties. For example, increasing competitive strength in the groups for which the condition *K_i_* > *N*_Σ_ is fulfilled leads to an increase in the community population, while for the other groups (for which *K_i_* < *N*_Σ_) the effect is the opposite. Since the responses (sensitivities) have different signs for different categories of groups, which we will term *functional group types*. Four functional types of groups can be identified: *super-dominant groups, dominant groups, sub-dominant groups and excluded groups*. With these, we can determine the structure of a multi-group community, as shown in the fig.2. Table 6 shows a detailed description of an example community composition, represented in terms of the functional group types. The relative proportions of the functionally different types of groups determines the composition of the community. fig.3 shows that as *D* increases, the relative proportions of the super-dominant and dominant groups decline, whereas the proportion of the excluded groups monotonically increases, and the proportion of the sub-dominant groups increases at small *D* then decreases.

**Fig.2.**
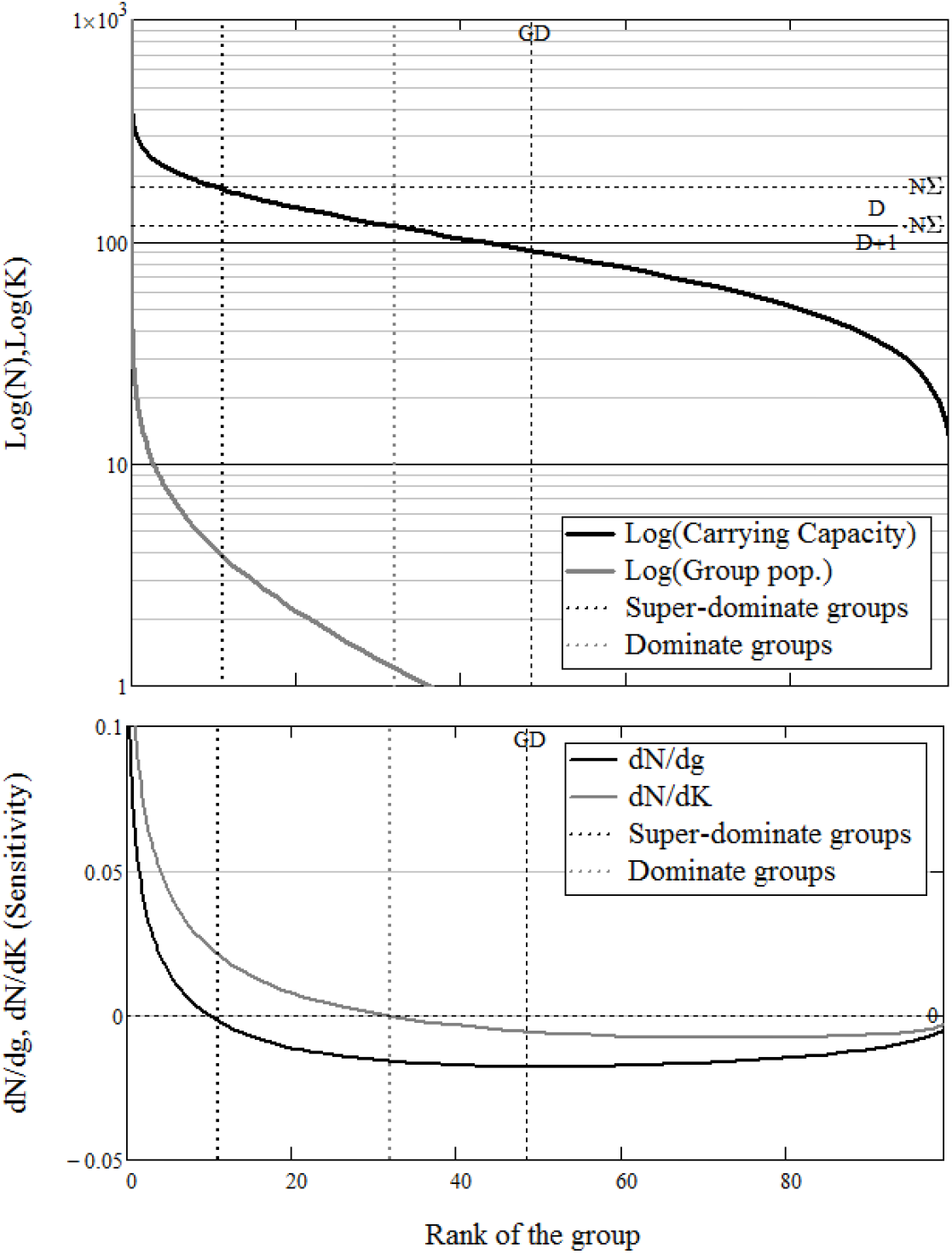
The structure of multi-group community. In upper panel, the rank distribution of singular currying capacities (bold black line) and the population size of the groups (grey line) for some multi-group community (with D=2, γ=const, K_i_ drawn from the beta distribution, m=100 groups) are shown. In corresponded bottom panel, the sensitivities of the total community population N_∑_ to competitive strength γ_i_ (black line) changes and to currying capacity K_i_ (grey line) changes are shown. Vertical lines from left to right show: Super-dominate group range (between zero and black dotted lines) for which dN_∑_/d*γ*_i_>0 and dN_∑_/dK_i_>0; Dominate group range (between black and grey dotted lines) for which dN_∑_/d*γ*_i_<0 and dN_∑_/dK_i_>0; Sub-dominate group range (between grey dotted and GD lines) for which dN_∑_/d*γ*_i_<0 and dN_∑_/dK_i_<0; Excluded group range (right to GD line).

**Fig.3.**
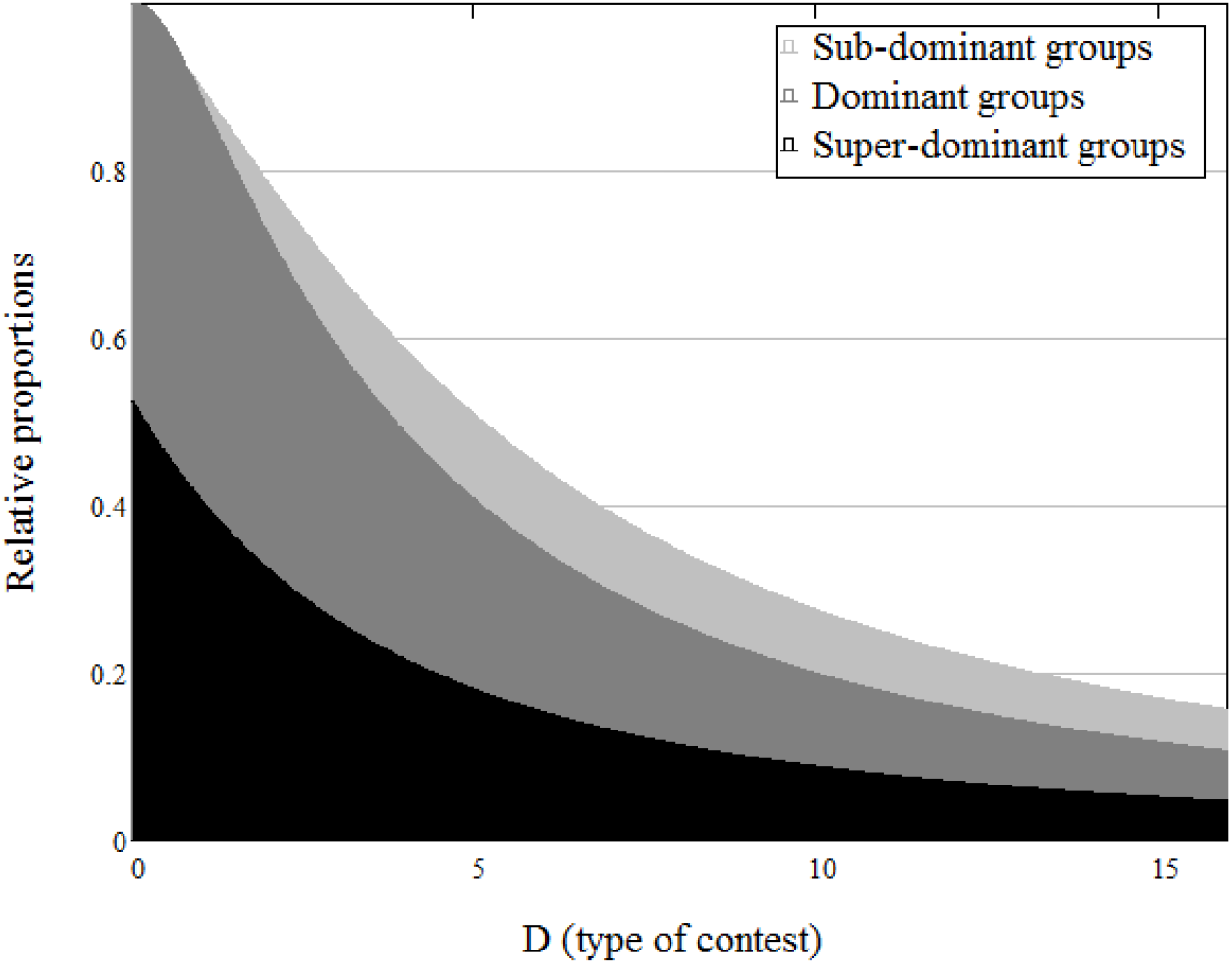
The composition of multi-group community in relation to contest type D, the relative proportion of super-dominant groups (black area), dominant groups (dark grey area), sub-dominate group (light grey area) and excluded groups (white area). The general trend of increasing the relative share of excluded groups as D increases demonstrates the principle of soft exclusion.

**Table 5.** The partial derivatives of i-th group population and the total multi-group community population with respect to (own and others) carrying capacities and to (own and others) competitive strengths.

| Partial derivative | Value of the derivative (sensitivity) | Sign of the derivative |
| --- | --- | --- |
| $\frac{\partial N_i}{\partial \gamma_i}$ | $\frac{D+1}{\gamma_i} N_i \left( 1 - \frac{N_i}{K_i} \right)$ | Always positive |
| $\frac{\partial N_i}{\partial \gamma_j}$ | $-\frac{D+1}{\gamma_j K_j} N_i N_j$ | Always negative |
| $\frac{\partial N_i}{\partial K_i}$ | $(D+1) \frac{N_i}{K_i} \left( 1 - \frac{D}{D+1} \frac{N_i}{K_i} \right)$ | Always positive |
| $\frac{\partial N_i}{\partial K_j}$ | $-D \left( \frac{N_j}{K_j^2} \right) N_i$ | Always negative |
| $\frac{\partial N_\Sigma}{\partial \gamma_i}$ | $\frac{D+1}{\gamma_i} N_i \left( 1 - \frac{N_\Sigma}{K_i} \right)$ | Positive for those groups for which the condition $K_i > N_\Sigma$ holds, otherwise negative |
| $\frac{\partial N_\Sigma}{\partial K_i}$ | $(D+1) \frac{N_i}{K_i} \left( 1 - \frac{D}{D+1} \frac{N_\Sigma}{K_i} \right)$ | Positive for those groups for which the condition $K_i > \frac{D}{D+1} N_\Sigma$ holds, otherwise negative |

**Table 6.** The structure of the multi-group community.

| Term | Definition | Conditions | Holds for groups for which |
| --- | --- | --- | --- |
| **Super-dominate group(s)** | Group(s) for which increasing their competitive strength leads to an increase in the total population of the multi-group community | $\frac{\partial N_\Sigma}{\partial \gamma_i} > 0$<br>$\frac{\partial N_\Sigma}{\partial K_i} > 0$ | $K_i > N_\Sigma$ |
| **Dominate group(s)** | Group(s) for which increasing their carrying capacity leads to an increase in the total population of the multi-group community | $\frac{\partial N_\Sigma}{\partial \gamma_i} < 0$<br>$\frac{\partial N_\Sigma}{\partial K_i} > 0$ | $N_\Sigma > K_i > \frac{D}{D+1} N_\Sigma$ |
| **Sub-dominate group(s)** | Group(s) for which increasing their competitive strength or their carrying capacity leads to a decrease in the total population of the multi-group community | $\frac{\partial N_\Sigma}{\partial \gamma_i} < 0$<br>$\frac{\partial N_\Sigma}{\partial K_i} < 0$ | $K_i < \frac{D}{D+1} N_\Sigma,$<br>$rank(K_i) < GD$ |
| **Excluded group(s)** | Group(s) that are not permanently present in the community, but may temporarily appear as "invaders" | $\frac{\partial N_\Sigma}{\partial \gamma_i} < 0$<br>$\frac{\partial N_\Sigma}{\partial K_i} < 0$ | $rank(K_i) > GD$ |

### 5. Fission-fusion and invasion-extinction dynamics

Above we considered the structures of ecological communities based on conspecific group composition, here we reconsider the structures of communities in the more familiar terms of multi-species populations. If we assume that groups belonging to the same species are ecologically equivalent (i.e., have the same set of density-independent parameters), then the population of the s-th species can be written as

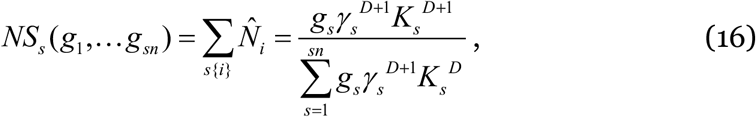

where *g_s_* is the number of groups within the s-th species population. Thus, the composition of a multispecies community can be represented by the configuration vector *g* (*g*,… *g_sn_*), where *sn* is the total number of species. It is reasonable to ask how the population structure of the community changes when the number of groups (constituting the species population) changes. To elaborate this, we introduce the fission-fusion (or divide-merge) operator as

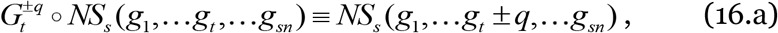

where the *+q* index refers to fission (i.e., increasing the number of groups within the t-th species population by *q*), and the *-q* index refers to fusion (i.e., reducing the number of groups by *q*). In the special case *q* = *-g_t_*, this can be called an extinction operator 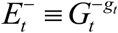, since in this case all groups belonging to the t-th species are removed from the community. Similarly, we can define an invasion operator as

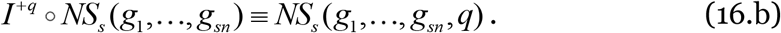

In addition, it is also useful to define the associated difference operators:

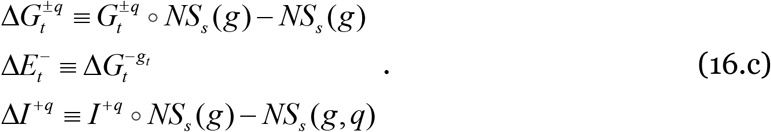

which calculate changes in species populations, where the indices *t* and *s* refer to focal and non-focal species, respectively. Table 7 provides a detailed description of how division-fusion within one species or an invasion/extinction of one of the species, affects all other species of the community, and hence the entire multi-species community. It is important to note that the community involved in the fission-fusion dynamic must be in a *transitive state*, since the equilibrium of the community depends on the group configuration *g*(*g*_l_,… *g_sn_*).

**Table 7.** The impact of fission-fusion and invasion-extinction events on species populations and the overall population of a multi-species community.

| Difference operator | Result | Interpretation of result |
| --- | --- | --- |
| $\Delta G_t^{+q} \circ N_s$ | $-\left(\frac{qN_t}{g_t K_t + qN_t}\right)N_s$ | Fission is always negative for populations of non-focal species |
| $\Delta G_t^{-q} \circ N_s$ | $\left(\frac{qN_t}{g_t K_t - qN_t}\right)N_s$ | Fusion is always positive for populations of non-focal species |
| $\Delta G_t^{+q} \circ N_t$ | $\left(\frac{q(K_t - N_t)}{g_t K_t + qN_t}\right)N_t$ | Fission is always positive for population of focal species |
| $\Delta G_t^{-q} \circ N_t$ | $-\left(\frac{q(K_t - N_t)}{g_t K_t - qN_t}\right)N_t$ | Fusion is always negative for population of focal species |
| $\Delta G_t^{+q} \circ N_\Sigma$ | $q\left(\frac{K_t - N_\Sigma}{g_t K_t + qN_t}\right)N_t$ | Fission is positive for the total population in the case of super-dominant focal species, and negative otherwise |
| $\Delta G_t^{-q} \circ N_\Sigma$ | $-q\left(\frac{K_t - N_\Sigma}{g_t K_t - qN_t}\right)N_t$ | Fusion is negative for the total population in the case of super-dominant focal species, and positive otherwise |
| $\Delta E_t^- \circ N_s$ | $\left(\frac{N_t}{K_t - N_t}\right)N_s$ | Extinction is always positive for populations of remaining species |
| $\Delta E_t^- \circ N_\Sigma$ | $-\left(\frac{K_t - N_\Sigma}{K_t - N_t}\right)N_t$ | Extinction is negative for the total population in the case of super-dominant species, and positive otherwise |
| $\Delta I_t^+ \circ N_s$ | $-\frac{N_t N_s}{K_t}$ | Invasion is always negative for populations of resident species |
| $\Delta I_t^+ \circ N_\Sigma$ | $\frac{(K_t - N_\Sigma)N_t}{K_t + N_t}$ | Invasion can be positive for the total population if the invasive species becomes a super-dominant, and negative otherwise |

The group-based approach allows for detailed tracking of the development of a multi-species community in terms of population size and structure change. Wide opportunities for theoretical and empirical research are presented using this tool, especially considering that the fission-fusion balance determines the rate of sympatric speciation. We next illustrate its practical usefulness with an applied problem that has proved difficult under the traditional approach.

### 6. Application: Calculating multi-species MSY for harvested populations

We consider a multi-group community consisting of *m* groups, in which *n* ≤ *m* groups can be harvested according to the catch rate vector *C_r_*(*c*_1_,…*c_n_*) (e.g., multi-stock fishery). We calculate how the total harvest *H*(*c*_1_,…,*c_n_*) depends on specific catch rates and find such values of the catch rate vector *C_MSY_*(*c*_1_,…*c_n_*) that maximize this function (i.e., the maximum sustainable yield: MSY). With additional harvesting, the population dynamics equation 6 can be rewritten as 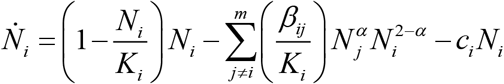, which for a fixed point can be rearranged as 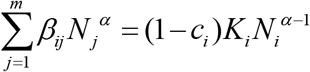. Comparing this expression with equation 10 it can be determined that the effect of harvesting is formally expressed as a reduction in singular carrying capacity by a factor of (1 – *c_i_*). Recall that here, for sake of simplicity, we use dimensionless catch rates, i.e., the dimensional rates are 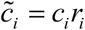, where *r_i_* are net growth rates. Next, using equation 13.a) we can find new equilibrium population sizes for each harvested (*k* ≤ *n*) and unharvested (*k* > *n*) group as

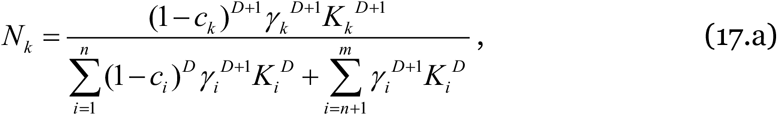

for harvested groups and

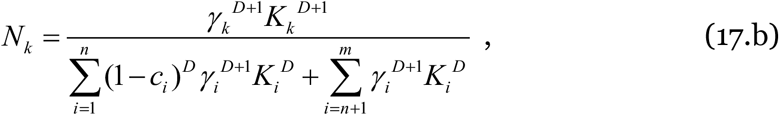

for unharvested groups. Note that harvesting always results in an increase in the equilibrium population size of unharvested groups (i.e., 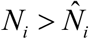). The total population size of the harvested community is

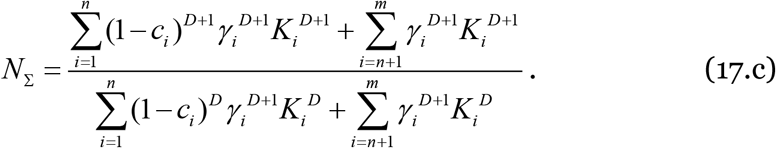

Note that for some values of the catch vector, harvesting may lead to an increase in the total population of the community.

The overall harvest 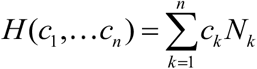 can be calculated as

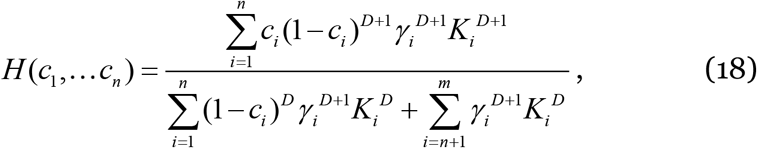

which in the multi-species terms becomes:

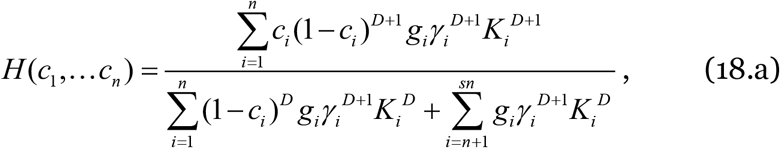

in which *c_i_* means species-specific catch rate and *n* is the number of harvesting species.

Using equation 18, we can immediately calculate multi-group (or multi-stock in the case of fishery) MSY in two extreme cases: (1) *D* → 0, which implies the situation of m completely independent (i.e., non-competing) groups and (2) *D* →∞, which describes the competitive exclusion situation (i.e., only one group survives in competition). In the first case, overall harvest is 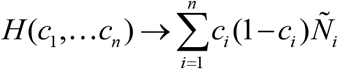 (where 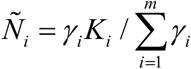 and the MSY catch vector is *C_MSY_* (*c*_1_,…*c_n_*) → (1 / 2,…,1 / 2).

In the second case, *H*(*c*_1_,…*c_n_*) → *c_k_* (1 – *c_k_*)*K_k_* (where *K_k_* is the singular carrying capacity of the group with *γ_i_ K_i_* the biggest value), and the MSY catch vector is *C_MSY_*(*c*_1_,…*c_k_*…*c_n_*)→(0,…1/2…0). The results in these extreme cases correspond to the well-known Schaefer model (Schaefer 1991), which is as expected since in these cases we are essentially dealing with single or several isolated groups.

For intermediate values of *D* at which, on the one hand, resource competition is significant, and, on the other hand, not strong enough to lead to the competitive exclusion of most groups, we can find the MSY catch vector by solving the equation ∀*H*(*c*_1_…*c_n_*) = 0, which can be reduced to the following system of equations:

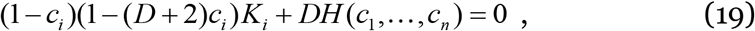

The first term of equation 19 is a simple quadratic form, and the second term is a fractional rational polynomial of degree *D* +1. It is not difficult to obtain a numerical solution for this system. It is also possible to obtain approximate solutions, which can be useful for obtaining “instantaneous” estimates of MSY rates and developing insight into natural resource management of multi-groups/multispecies communities. As an example, we first consider the case with *n* = 1 (i.e., only one group is harvested) and examine how the position of the focal group within a community affects the community-level MSY catch rate. In this case, it can be shown (Appendix A) that the solutions of the system of equations 19 lie within the interval 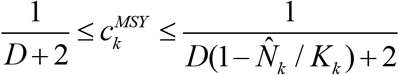. Fig. 4 shows the structure of a multi-group community (i.e., structured into functional group types) for various values of *D* with corresponding values of 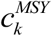. If desired, solutions can also be found for the case *n* = 2, which includes 10 subcases (i.e., the number of unique pairs from 4 categories). Extending beyond, for cases *n* ≥ 3, the number of subcases grows exponentially, which makes it difficult to write solutions. On the other hand, approximate solutions to this system can be represented by the combination of two structural parts: (1) solutions for super-dominant group for which *DH*(*c*_1_,…*c_n_*) ≈ *Dc_i_*(1 – *c_i_*)*K_i_* (meaning that overall harvest is mostly defined by one group) and (2) solutions for dominant and subdominant groups for which *DH*(*c*_1_,…*c_n_*) ≈ *const* (meaning that the contribution to overall harvest of each group is relatively small). The MSY catch rate for super-dominant group is close to 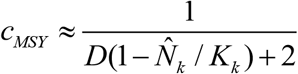. In turn, for the dominant and subdominant groups, equation 19 reduces to a quadratic equation that has no real roots if *K_i_* < *K_low_*, where 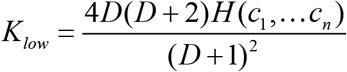, and has two real roots:

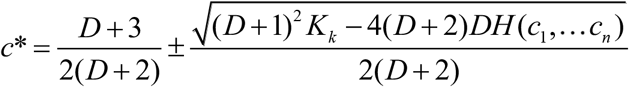 otherwise. The values of the smaller root are in the range 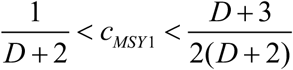, and the values of the larger root are in the range 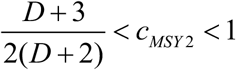.

**Fig.4.**
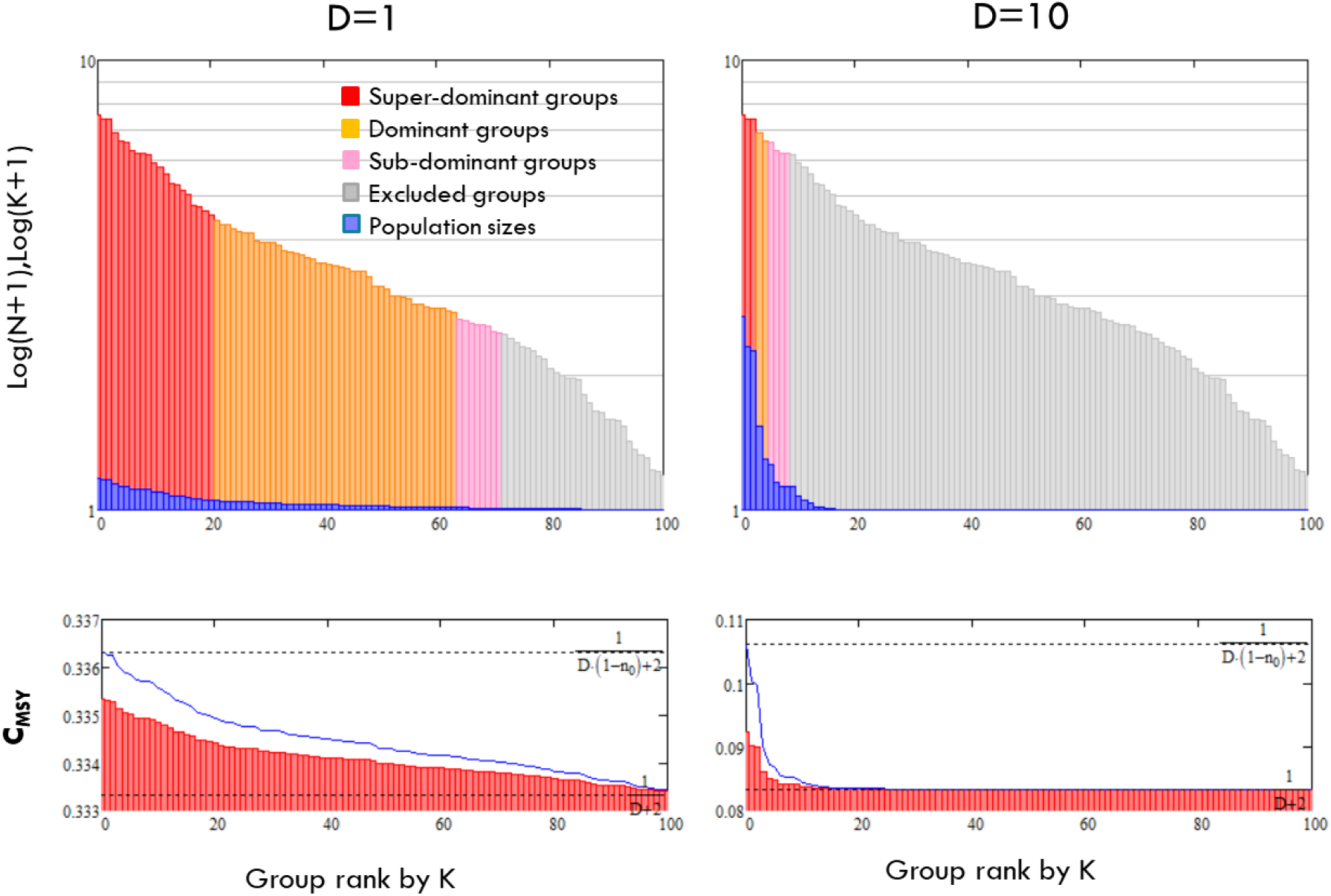
The structure of multi-group community and MSY for single harvesting group at different levels of D. In the top panels, singular currying capacities of

These implicit solutions allow some generalised conclusions about multi-species MSY:

1. Multi-species MSY is not necessarily unique, in some cases, there may be several combinations of catch rates that meet MSY conditions (following the tradeoff between relatively small exploitation of many stocks and heavy exploitation of a few stocks).
2. MSY of super-dominant groups is weakly dependent on the harvesting of other groups, which means that the standard single-species MSY approach remains valid if exploited stocks are super-dominant in the community.
3. Groups with a singular carrying capacity below the critical *K_low_* cannot be sustainably harvested.

Regarding the latter, note that a low singular carrying capacity does not mean a low abundance of this species, as the total species population may consist of many conspecific groups. However, an attempt to manage such a stock based on standard (single species) MSY estimates can lead to its rapid depletion, since such species are already “overexploited” by other species within the community.

Essentially, the management of multi-species communities requires knowledge of the position of these species within the community. In some cases (i.e., with super-dominant species) no significant changes in existing practice are required, in other cases (dominant species) it is necessary to consider the existence of several solutions, finally in the case of subdominant species the approach based on single-species MSY needs to be radically reconsidered (for example that is quite likely the case with many elasmobranch species).

## 7. Discussion

In this section, we discuss the relative position of the group-based approach and the multi-group theory among other approaches and theories, noting essential differences and focusing on similarities to highlight their intermediate nature. Table 8 contrasts the group-based approach with the most common alternatives: the species-centric and trait-based approaches.

**Table 8.** Features of Species-centric, Group-based, and Trait-based approaches to modelling ecological communities.

|  | **Species-centric (classical) approach** | **Group-based approach** | **Trait-based approach** |
| --- | --- | --- | --- |
| **Conceptual hierarchy of ecological community** | Individual<br>Species<br>Multi-species community | Individual<br>Conspecific Group<br>Species<br>Multi-group/species community | Individual<br>Community of individuals with different traits |
| **Basic entities** | Species | Conspecific groups | Individuals |
| **Key interactions** | Between species populations | Between conspecific groups | Between individuals, regardless species affiliations |
| **Fitness defined as** | Individual and species specific | Inclusive and group/species specific | Individual and trait specific |
| **Community structure is described in terms of** | Relative species abundances | The impact that a group has on other groups and on the community | Trait distributions |
| **Modelling by** | System of Lotka-Volterra equations | System of resource sharing and resource-based population dynamics equations | System of trait-dependent growth and trade-off between key traits equations |
| **The coexistence of species is ensured by** | Fine-tuning the species parameters or introducing “stabilizing/equalizing mechanisms” | Self-regulation of the sizes of conspecific groups | Tuning the trade-off parameters |
| **The coexistence is** | Evolutionarily unstable | Evolutionarily stable | Uncertain |

Despite the differences, there is a direct formal connection between the classical (species-centric) and group-based approach, as established by the equation 6, which shows that the former is a special case of the latter under the *α* = 1 condition. In other words, both approaches are equivalent in the absence of group behaviour, otherwise the group-based perspective can be seen as an analytical extension of the Lotka-Volterra competition model beyond the standard type of interaction term.

Connection with the R* resource-ratio theory (Tilman, 1999, 2014) can be formally established by calculating the amount of resources needed to support a minimum viable population size *MVP_i_* as 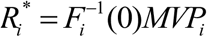 (equation 2), these values can be a good proxy for estimating singular carrying capacities (equation 3), which means that these values in some degree define species (group) abundances. However, within the group-based approach, these 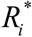 quantities do not generally determine species (group) coexistence, except in the classic *α* = 1 case. In other words, if a species cannot ensure its existence in a community due to limited resources, this does not mean that this species cannot coexist in this community, for example, when resource constraints are relaxed.

In contrast to modern coexistence theory (Chesson, 2000; HilleRisLambers et. al. 2004), the group-based approach shifts the focus from intra-inter-species to intra-inter-group relationships. This shift allows us to find parsimonious (without the involvement of additional equalizing and stabilizing mechanisms) solution for the coexistence puzzle, namely, depending on the obtained resource, the group naturally regulates its size in such a way that it balances intra-group and inter-group competition. From the point of view of modern co-existence theory, group behaviour can be seen as a specific combination of stabilizing and equalizing mechanisms, and the ubiquitous presence of groups as a natural solution to the coexistence problem. Significantly, the group-based approach considers the principle of competitive exclusion not as a condition for extinction, but rather as a precondition for the emergence of group behaviour.

Parallels can be drawn with the niche construction theory (Odling-Smee et.al. 2013): individuals entering group relations (e.g., preferring to interact with some individuals and avoiding others), directly change their biotic environment and construct an evolutionary niche (Trappes, 2021). The construction of such an evolutionary niche allows species to coexist stably. In this context, multi-group theory can be viewed as a specific model of biodiversity within the framework of the extended evolutionary synthesis (Laland, 2014).

There is a long-standing controversy about the way ecological processes determine relative species abundance (McGill, 2003; Connolly *et al*. 2014), in which modern niche theory (Letten, 2017) and unified neutral theory (Hubbell, 2005) take opposite sides. Without claiming final resolution, multi-group theory can be a bridge between the two, as it allows the tracking of changes in relative species abundance as neutrality conditions are gradually introduced (i.e., *Var*(*C*) → 0 and *Var* (*γ*) → 0). The results of this and some other transitions to the limits can be seen in the table 4. In the case of a lognormal distribution of competitive ability, equations 14 reveals the relationship between relative group abundance, the diversity index, and the community population. It can be noted that the transition to neutrality (i.e., *GD* → *m*) is not universal and depends on the type of competition (i.e., qualitative vs. quantitative) in the community. In a community with strong quantitative competition (*D*» 1), small variations in competitive abilities can significantly reduce diversity, but in a community far from the competitive exclusion limit (*D*≪ 1), even large variations do not greatly affect diversity and may not violate neutrality. Thus, testing the predictions of the neutral theory requires considering the competition milieu, otherwise the results may be inconsistent and/or dependent on specific communities.

Although the trait-based and group-based approaches diverge on some points (e.g., the focus on absolute traits and individual fitness vs. a focus on relative traits and inclusive fitness), there are also significant overlaps in some other themes such as interaction milieu, performance currency and environmental gradients (McGill et.al. 2006). In multigroup theory, the interaction milieu is quantified by the parameter alpha *α* (or *D* in the case of *α* < 1), which can be interpreted in two ways: theoretically as the relation between quantitative and qualitative factors of resource competition and empirically as the order of diversity *q* in the Hill number. Also, performance currency can be represented by the energy content of a resource, but this currency concept is limited to the case of zero-sum resource competition, in which the performance of each group can be uniquely determined in terms of the obtained resources. In other cases (including the classical Lotka-Volterra model), performance cannot be unambiguously determined (for example, species A performs better than B, B is better than C, but C is better than A), which means the impossibility of one universal performance currency or (following this analogy) the need for a multi-currency system to describe ecosystems. Within the multigroup theory, environmental gradients (at the macro scale) can be represented by an ∇*R* gradient of available resources and an ∇*α* gradient. The theory can predict the response of community structure to each of these gradients. However, the question of a possible correlation between these gradients remains open to empirical testing.

The multi-group theory of biodiversity overlaps with the domains of other community theories in many ways, providing an opportunity for synthesis and unification. Importantly, it can bridge the gap between inefficient theories (i.e., based on a large number of free parameters) and efficient ones (i.e., based on a few first principles).

Like any other approach, the group-based one has its advantages and limitations, which can be divided into two areas: empirical and analytical. Although the presence of conspecific groups within species is ubiquitous, empirical data for this remains sparse with the focuses more on species rather than group affiliation. Modern monitoring methods and the growing interest in intra-species trait variation (Violle, C. *et al*. 2012) should eventually eliminate this discrepancy and allow direct verification of the theory. Although the group approach has found an exact analytical solution to the problem of species coexistence in the case of zero-sum resource competition, in other cases it faces the same analytical difficulties as the classical models of competition. However, the presence of a smooth analytical transition between cases can significantly simplify the search for numerical solutions in non-zero-sum competition cases.

Ultimately, the main criterion for the significance of a theory is practical application. The group-based approach offers a method for finding the MSY in the case of a multi-species community, which is much simpler than those currently used, since it relies more on analytical results than on numerical simulations. In addition, the equation 18.a can be directly used in economic models of natural resource management instead of multiple instances of the Schafer model used for a single-species MSY.

## 8. Acknowledgments

This work was supported partially by Department of Agriculture, Food and the Marine (DAFM) (FishKOSM 487 project, DAFM reference 15/S/744).

## 9. Conflict of interest

Authors declare no conflict of interests.

## Notes

### Competing Interest Statement

The authors have declared no competing interest.

## References

Andersen, K. H., Jacobsen, N.S. and Farnsworth, K.D. (2016). The theoretical foundations for size spectrum models of fish communities. Canadian Journal of Fisheries and Aquatic Sciences. 73(4): 575–588.

Carvalho, J. C. & Cardoso, P. (2020). Decomposing the Causes for Niche Differentiation Between Species Using Hypervolumes. Front. Ecol. Evol. 8, 1–7.

Chesson, P. (2000). Mechanisms of maintenance of species diversity. Annual Review of Ecology and Systematics. 31: 343–366.

Connolly, S. R. et al. (2014). Commonness and rarity in the marine biosphere. Proc. Natl. Acad. Sci. U. S. A. 111, 8524– 8529.

Crowley, PH, Hwan Baik K. (2010). Variable valuations and voluntarism under group selection: An evolutionary public goods game. J Theor Biol 265:238–244.

Gause, G.F. (1934). The struggle for existence. Baltimore: Williams & Wilkins.

Gavrilets, S. (2012). Human origins and the transition from promiscuity to pair-bonding. Proc Natl Acad Sci USA 109:9923–9928.

Gavrilets, S. (2012). On the evolutionary origins of the egalitarian syndrome. Proceedings of the National Academy of Sciences, 109(35), 14069–14074.

Haldane, J. B. S. (1932). The causes of evolution. London: Longmans, Green & Co.; New York: Harper Brothers.

Hamilton, W.D. (1963). The Evolution of Altruistic Behavior. The American Naturalist, 97, 354–356.

Hardin, G. (1960). The competitive exclusion principle. Science. 131 (3409): 1292–129x.

Hawkes K, Rogers AR, Charnov EL (1995) The male’s dilemma: Increased offspring production is more paternity to steal. Evol Ecol 9:662–67x.

Hill, M. O. (1973). Diversity and evenness: a unifying notation and its consequences. Ecology. 54: 427–432.

HilleRisLambers, J., Adler, P. B., Harpole, W. S., Levine, J. M., & Mayfield, M. M. (2012). Rethinking community assembly through the lens of coexistence theory. Annual Review of Ecology, Evolution and Systematics, 43(227).

Hirsch, M. W. (1988). Systems of differential equations which are competitive or cooperative: III. Competing species. Non-linearity, 1, 51–71.

Hubbell, S. P. (2005). Neutral theory in community ecology and the hypothesis of functional equivalence. Funct. Ecol. 19, 166–172.

Hubbell, S.P. (2001). The Unified Neutral Theory of Biodiversity and Biogeography. Princeton University Press. ISBN 978069102128x.

Hutchinson, G,E. (1961). The paradox of the plankton. American Naturalist. 95: 137–145.

Jost, L. (2006). Entropy and diversity. Oikos. 113: 363–375.

Kiørboe, T., Visser, A., Andersen, K. H. & Browman, H. A. (2018). trait-based approach to ocean ecology. ICES J. Mar. Sci. 75, 1849–1863.

Kokko H, Morrell LJ (2005). Mate guarding, male attractiveness, and paternity under social monogamy. Behav Ecol 16:724–731.

Konrad, K. (2009). Strategy and Dynamics in Contests (Oxford Univ Press, Oxford).

Krause, J., & Ruxton, G. D. (2002). Living in groups. Oxford: Oxford University Press.

Laland KN, Uller T, Feldman M, Sterelny K, Müller GB, Moczek A, Jablonka E et al (2014). Does evolutionary theory need a rethink? Nature 514(7521):161–164.

Letten, A. D., Ke, P. J. & Fukami, T. (2017). Linking modern coexistence theory and contemporary niche theory. Ecol. Monogr. 87, 161–177.

MacArthur, R and R Levins. (1967). The Limiting Similarity, Convergence, and Divergence of Coexisting Species. The American Naturalist 101(921): 377–385.

Marquet, P. a. et al. (2014). On Theory in Ecology. Bioscience 64, 701–710.

May, R. (2019). Stability and Complexity in Model Ecosystems. Princeton: Princeton University Press.

McGill, B. J. (2003). A test of the unified neutral theory of biodiversity. Nature 422, 881–885.

McGill, B. J., Enquist, B. J., Weiher, E. & Westoby, M. (2006). Rebuilding community ecology from functional traits. Trends Ecol. Evol. 21, 178–185.

Odling-Smee, F. J., Laland, K. N., & Feldman, M. W. (2013). Niche construction. In Niche Construction. Princeton university press.

Pocheville, A. (2015). The Ecological Niche: History and Recent Controversies. In Heams, Thomas; Huneman, Philippe; Lecointre, Guillaume; et al. Handbook of Evolutionary Thinking in the Sciences. Dordrecht: Springer. pp. 547– 586. ISBN 978-94-017-9014-x.

Reeve, HK, Hölldobler B (2007). The emergence of a superorganism through intergroup competition. Proc Natl Acad Sci USA 104:9736–9740.

Sadykov, A. (2011) Conspecific Community Dynamics Models and their applications, Series of dissertations, Unipab ISSN 1501-7710.

Schaefer, M.B. (1991). Some aspects of the dynamics of populations important to the management of the commercial Marine fisheries. Bltn Mathcal Biology 53, 253–279.

Schelling, TC (1978) Micromotives and Macrobehavior (Norton, New York).

Smale, S. (1976). On the Differential Equations of Species in Competition. J. Math Biol., 3, 5–x.

Tilman, D. (1999). The ecological consequences of changes in biodiversity: Perspectives. Ecology 80 (5): 1455–74.

Tilman, D., Isbell, F., & Cowles, J. M. (2014). Biodiversity and ecosystem functioning. Annual review of ecology, evolution, and systematics, 45, 471–493.

Townsend C, Begon M, Harper JL. (2008). Essentials of Ecology. John Wiley & Sons; 3rd Edition. ISBN-10: 1405156589.

Trappes, R. (2021). Defining the niche for niche construction: evolutionary and ecological niches. Biol Philos 36, 31.

Tullock, G. (1980). Toward a Theory of the Rent-Seeking Society, eds Buchanan JM, Tollison RD, Tullock G (Texas A&M University, College Station, TX), pp 97–112.

Vinković, D. Kirman, A. (2006). A Physical Analogue of the Schelling Model. Proceedings of the National Academy of Sciences 103 (51): 19261–65.

Violle, C. et al. (2012). The return of the variance: Intraspecific variability in community ecology. Trends Ecol. Evol. 27, 244–252.

